# The vacuolar protein 8 (Vac8) homolog in *Cryptococcus neoformans* plays conserved and unique roles in vacuolar and cellular morphology, impacting important stress responses and virulence traits

**DOI:** 10.1101/2025.05.22.655601

**Authors:** Peter V. Stuckey, Julia Marine, Meghan Figueras, Aliyah Collins, Felipe H. Santiago-Tirado

## Abstract

Functionally similar to a plant vacuole or a mammalian lysosome, the fungal vacuole plays a vital role in many cellular processes. Most studies of the vacuole have been performed in the nonpathogenic yeast *Saccharomyces cerevisiae*; however, recently in pathogenic fungi, the vacuole has been implicated in host invasion in both plants and mammals highlighting an important role for the vacuole in pathogenesis. Here, we report that deletion of *Cryptococcus neoformans* vacuolar protein 8 (*VAC8*) results in a fragmented vacuole morphology, impairment of vacuolar fusion, and inability to form titan cells. Additionally, absence of Vac8 results in defective growth at high temperature and in the presence of caffeine, suggesting a defect in cell wall signaling. Interestingly, despite aberrant vacuole morphology, *vac8Δ* is slightly more resistant to fluconazole treatment, and displays increased resistance to hydrogen peroxide, suggesting the irregular vacuole morphology does not impair vacuole function. Like *S. cerevisiae* Vac8, *C. neoformans* Vac8 is comprised of armadillo repeat regions which form alpha helices that fold to form a superhelix allowing for increased protein-protein interaction. Many of the known binding partners of *S. cerevisiae* Vac8 are not present in the *C. neoformans* genome, suggesting novel functions for Vac8 in this fungus. Notably, deletion of *VAC8* affected some virulence traits, providing support to targeting the fungal vacuole as a potential therapeutic intervention.

## INTRODUCTION

The incidence of serious and/or invasive fungal infections has been increasing in the last few decades, with overall mortality rates >20%. (*1*) Despite this, there have been few advances in available antifungal treatment options with most targeting cell wall components or ergosterol, a cholesterol-like lipid found only in fungi. (*2*) Additionally, these medications are complicated by off target toxicity, restrictive formulations, limited activity, restrictive availability, and the emergence of antifungal resistance.(*3, 4*) Therefore, morbidity and mortality rates from systemic fungal infections remain incredibly high despite a decrease in the HIV/AIDS populations, which are the most susceptible to these infections.(*5*) This highlights the need for novel antifungal therapies and the identification of potential new drug targets.

The fungal vacuole is a dynamic organelle that reacts to a variety of environmental stresses and changes its morphology accordingly. Functionally similar to a plant vacuole or a mammalian lysosome, the fungal vacuole plays a vital role in degradation of intra- and extracellular substrates, recycling of nutrients, metabolite storage, and detoxification.(*6, 7*) To perform this diverse set of functions, the vacuolar membrane is constantly undergoing coordinated fusion and fission events to respond to changes in the environment. Most studies of fungal vacuoles have been performed in the nonpathogenic yeast *Saccharomyces cerevisiae,* where multiple genes involved in vacuolar functions have been characterized (known as *VAC* genes). In *S. cerevisiae*, *VAC8* (*ScVAC8*) has been extensively studied given that it encodes the only Armadillo (ARM) repeat-containing protein and is involved in many cellular roles including vacuole fusion, vacuole inheritance, and autophagosome formation, among others.(*8*) Therefore much of what we assume to know about the vacuole in pathogenic fungi comes from *S. cerevisiae* despite there being large genetic differences. We do know that at least some of these vacuolar functions are critical for some pathogenic species to survive and invade mammalian hosts and cause disease. (*9–11*)

In human fungal pathogens, the role of the vacuole in basic cell biology and during infection is largely unexplored but recently has been shown to play a critical role in supporting host invasion in mammals. *Candida albicans* and *C. neoformans* fungal mutants with greatly impaired vacuole function do not cause lethal disease in mouse models of disseminated disease.(*9–11*) In *C. neoformans*, deletion of a vesicular proton pump, VPH1, leads to defects in major virulence factors including capsule production, reduction in thermotolerance, laccase function, and urease secretion.(*12*) This lead to complete loss of virulence in a mouse model of disease. Likewise, in other fungal pathogens like *Nakaseomyces glabratus* (formerly *C. glabrata*) and *Histoplasma capsulatum*, normal vacuolar function has been implicated in the ability to cause disease in mouse models.(*13, 14*)

Particularly for *C. neoformans*, the interactions with host immune cells are critical for determining the outcome of the infection, and can play a key role in the dissemination to the central nervous system.(*15, 16*) For example, clinical, *in vivo*, and *in vitro* data have demonstrated a strong correlation between phagocytosis of *C. neoformans* by macrophages and a negative outcome for patients.(*17, 18*) Isolates with high rates of *in vitro* phagocytosis and high intracellular proliferation correlated with patient death (despite treatment they all died at 3 months).(*18*) Similarly increased rates of *in vitro* phagocytosis correlated with high fungal burden on the brain, which increases the risk of mortality.(*17, 18*) Along these lines, our lab previously sought to screen a single gene deletion collection of *C. neoformans* for mutants that display altered phagocytosis *in vitro* by THP-1 human macrophages.(*19*) One of the identified mutants, missing the gene *PFA4* that encodes an S-acyl transferase that catalyzes lipid modifications of proteins, displayed severe defects in a wide variety of cell stressors and was completely avirulent in an intranasal mouse infection model.(*12, 19*) In an effort to mechanistically explain the phenotypes, we used click chemistry to identify the specific proteins modified by Pfa4, in the process determining the first “palmitoylome” of any fungal pathogen.(*19*) One of the substrates was the protein encoded by CNAG_00354, the homolog of *ScVAC8*, which is known to be palmitoylated in both *S. cerevisiae* and *Toxoplasma* species.(*20, 21*) Given the pleiotropic effects exhibited by the *pfa4Δ* mutant and the known involvement of *ScVAC8* in vacuolar functions, we wondered if *VAC8* would have similar functions in *C. neoformans*. If true, vacuolar dysfunction might be one of the factors contributing to the lack of virulence in *pfa4Δ*.

Here, we report that indeed the *pfa4Δ* mutant shows vacuole defects similar to those shown by deletion of just *VAC8* (*vac8Δ*). ScVac8 is palmitoylated and myristoylated, which is required for targeting Vac8 to the vacuole membranes in *S. cerevisiae*.(*22*) Given that *C. neoformans* Vac8 also contains the same lipid-modified residues as ScVac8, it would be expected that in the *pfa4Δ* mutant, Vac8 would be mis- localized, affecting vacuolar function. To carefully examine the effects of *vac8Δ*, we visualized vacuole morphology in wildtype (*23*) and *vac8Δ C. neoformans* strains, finding that deletion of *VAC8* leads to aberrant and fragmented vacuolar structure. Additionally, the *vac8Δ* strain displays increased sensitivity to growth at high temperature and impaired growth in the presence of caffeine, suggesting a defect in cell wall stress signaling. Notably, *vac8Δ* shows increased resistance to fluconazole (FLC), but no change to amphotericin B. Interestingly, *vac8Δ* strains also show increased resistance to high levels of oxidative stress (H_2_O_2_) compared to WT strains. The pleiotropic phenotypes of *vac8Δ* are not surprising given the multiple binding partners Vac8 might have. The multiple ARM repeats allow Vac8 to mediate multiple processes, some of which are important for virulence, making Vac8 (or vacuolar functions) an optimal target for possible therapeutic intervention.

## MATERIALS AND METHODS

### Strains, cell lines, and growth conditions

Fungal cells were thawed from -80°C stocks every 6 months and struck onto YPD plates allowing for 48 hours of outgrowth at 30°^C^ prior to storage at 4°^C^ for up to a month before being replated. Cells were grown in YPD liquid media overnight at 30°C with shaking (225 rpm). Cultures were then diluted back to OD_600_ of 0.2 and allowed to double until OD_600_ reached 0.5-1.0, unless otherwise stated.

### In silico sequence analysis

Sequences of proteins analyzed in this paper were obtained from NCBI (www.ncbi.nlm.nih.gov) or from FungiDB (fungidb.org). To search for Vac8 homologues the ScVAC8 and *C. neoformans VAC8* sequences were BLASTed against other common fungal pathogens. Selected proteins representing Basidiomycota, Ascomycota, and Mucorales were utilized to create a phylogenetic tree. Phylogenetic analysis was performed using maximum likelihood method in MEGA11 software and confidence intervals were determined from 1000 bootstraps with bootstrap scores shown at all nodes. The tree shown is the bootstrap consensus tree.

Protein domain analysis was performed using Interpro and Pfam database. Protein structures are predicted and visualized using Alphafold3 for *C. neoformans*, *S. cerevisiae*, and *C. albicans*.

Palmitoylation and myristylation was determined using palmitoylation prediction software GPS Palm (*24*) and myristylation was determined using software (*25*).

### Drug susceptibility assays

The MIC of Amphotericin B, and Fluconazole was determined on RPMI plates with Epsilometer test strips (E-test strips; Liofilchem Thermo). 5 × 10^4^ log phase cells in 200 uL PBS were plated on RPMI agar plates, spread evenly across the plate and allowed to dry before application of an E-test strip. Plates were incubated at 37° C and 5% CO_2_ and imaged after 72 hours.

### Stress Plate Phenotyping

Strains were grown overnight in YPD at 30°C with shaking (225 rpm) to an OD600 of 0.5-1.0. Cells were washed with PBS and the concentration was adjusted to 10^7^ cells/mL in PBS. Cultures were then serially diluted 10-fold in PBS and spotted (5 uL) onto solid YPD ager medium and incubated at 30° C, unless otherwise noted, for 3 days and imaged daily. The response of various *vac8Δ* mutants was tested under the following conditions: 1M sodium chloride (NaCl); 0.75 mg/mL caffeine; 0.3% sodium dodecyl sulfate (SDS); 2mM and 4 mM hydrogen peroxide (H_2_O_2_); 2 mM nitrosative stress (NaNO_2_); YPD, and RPMI plates were also tested at 37° C.

### Fungal Genomic Manipulation

The commercially available CNAG_00354 single gene deletion mutant was obtained from the *Cryptococcus* deletion collection (Fungal Genetics Stock Center). To complement this strain the coding sequence of *VAC8* was amplified via PCR with primers containing restriction digest sites for FseI and NheI. The PCR product was then digested with FseI and NheI in cutsmart buffer and ligated into plasmid pDC-mRuby3- neo which is derived from pGWSK7 (mRuby). (*26*) This plasmid contains a histone 3 promoter and mRuby3 tag at the C-terminus. The entire cassette including neomycin resistance was amplified via PCR using primers with 50 bp homology to the safe-haven 2 (SH2) region. Cas9 was amplified from pBMH2403_CnoCas9, and the guide was amplified from a derivative of pBHM2329. (*27*) DNA was then electroporated into the *vac8Δ* mutant using standard electroporation settings. Briefly, cells were grown overnight in YPD, then diluted back to OD600: 0.2 and allowed to grow until mid-log phase. Cells were than washed twice with ice cold diH_2_O and resuspended in 10 mL electroporation buffer (EB) with DTT. Cells were then pelleted are resuspended in 250 uL EB without DTT and electroporated in a 2 m gap cuvette with 45 uL cells, 500 ng Cas9, 350 ng sgRNA, and 2000 ng repair construct, at 2000V, 25 uF, and 200 Ω. Cells were immediately supplemented with YPD and incubated at 30° C with rotation for 2 hours prior to plating on YPD+Neo plates and incubated at 30° C for 72 hours. Complementation was confirmed by amplifying the entire SH2 region and sequencing.

### Titan cell induction and quantification

Cells are grown overnight in 5 mL YNB at 30° C with shaking (225 rpm). Log phase cells are harvested and washed 2X with PBS and counted using a Bio-Rad TC10 cell counter. Cell density is then adjusted to 1000 cells/mL in 10mL of 10% heat-inactivated fetal bovine serum (FBS) in PBS in a T25. Flasks are placed in 37° C with 5% CO2 and incubated for 48 hours before being washed and imaged. Cell size, capsule size, and total cell size were quantified manually in FIJI using the average of four measurements per cell.

### Vacuole induction and FM4-64 staining

To visualize the vacuole of *C. neoformans* strains cells were stained with FM4-64 and calcofluor white (CFW) to highlight the cell wall. Briefly, cells were grown under specified conditions, washed 2X with PBS and resuspended in 100 μL of PBS. 2 µl of FM4-64 (32 µM final concentration) is added and incubated at 30° C for 30 min with rotation. Cells are then washed 2X with PBS and resuspended in 50 μL of PBS and imaged under a Zeiss Axio Observer 7, with an Axiocam 506 mono camera and a FM4-64 long-pass filter set, using 100X/1.4 plan-apo oil objective. To induce the large vacuole and test for vacuole fusion, cells were grown overnight in YPD+ 1 M sorbitol at 37° C, then washed and grown in YPD + 1 M sorbitol at 37° C for 4 hours prior to staining and imaging and done previously. (*28*)

### Mouse virulence study

Female A/J mice 5-6 weeks old (Jackson Laboratory) were used in this study. All animals had free access to food and water. All testing and treatment of the mice in the study was performed in accordance with the University of Notre Dame and its Animal Care and Use Committee under the protocol number 23-05-7918. Briefly, fungal strains were cultured overnight in YPD to mid log phase, collected, washed and diluted to 1.25x10^6^ cells/mL in DBPS. Mice were intranasally infected with 40 μL of the cell solution (5x10^4^ cells) or 40 μL of DPBS under anesthesia. Animals were monitored daily and sacrificed at set time points of 3-, 7-, and 14-days post infection, or when they lost >20% of their peak weight. To quantify organ fungal burden the lungs, brain, and spleen were homogenized using a VWR 200 homogenizer and plated on YPD+AMP agar. After 48 hours CFUs were enumerated.

### Histology

A/J female mice (The Jackson Laboratories) were infected via intranasal route as described above with 5 x 10^4^ CFU of either WT (KN99), the *vac8Δ* mutant strain, *vac8Δ*:VAC8, or with sterile DPBS. Mice were sacrificed as predetermined time points (3, 7, and 14 days post inoculation, and at TOD) by isoflurane overdose followed by cervical dislocation. Lungs were perfused with 10% neutralized formalin and subsequently paraffin embedded, sectioned, mounted, and stained with H&E by the University of Notre Dame Histology Core facility. H&E-stained slides were imaged at 10X and 40X using an inverted Zeiss Axio Observer 7.

## RESULTS

### Deletion of both *PFA4* and CNAG_00354 causes vacuolar fragmentation

Since CNAG_00354 is a homolog of *ScVAC8*, a key vacuole regulatory protein involved in vacuolar inheritance and fusion, nuclear-vacuole junctions, and autophagy, among others, (*29*) we wondered if CNAG_00354 was also involved in vacuolar function. We used the lipophilic dye FM4-64 to visualize the vacuole of WT, *pfa4Δ*, and cnag_00354Δ grown under standard media conditions (YPD at 30°C). We saw distinct but overlapping phenotypes (**Figure 1A**). In WT cells there are usually 1 – 2 big vacuoles, with few tubules or smaller vesicles visible. In the cnag_00354Δ mutant the vacuole was completely fragmented into small vesicles without tubules, while in the *pfa4Δ* there was an intermediate phenotype. Notably, there was no obvious vacuole inheritance defect, as we could see vacuoles present in bud and daughter cells, suggesting that CNAG_00354 is either not performing that function as in *S. cerevisiae* or there are redundant pathways in *C. neoformans* to ensure vacuole inheritance.

**Figure 1:**
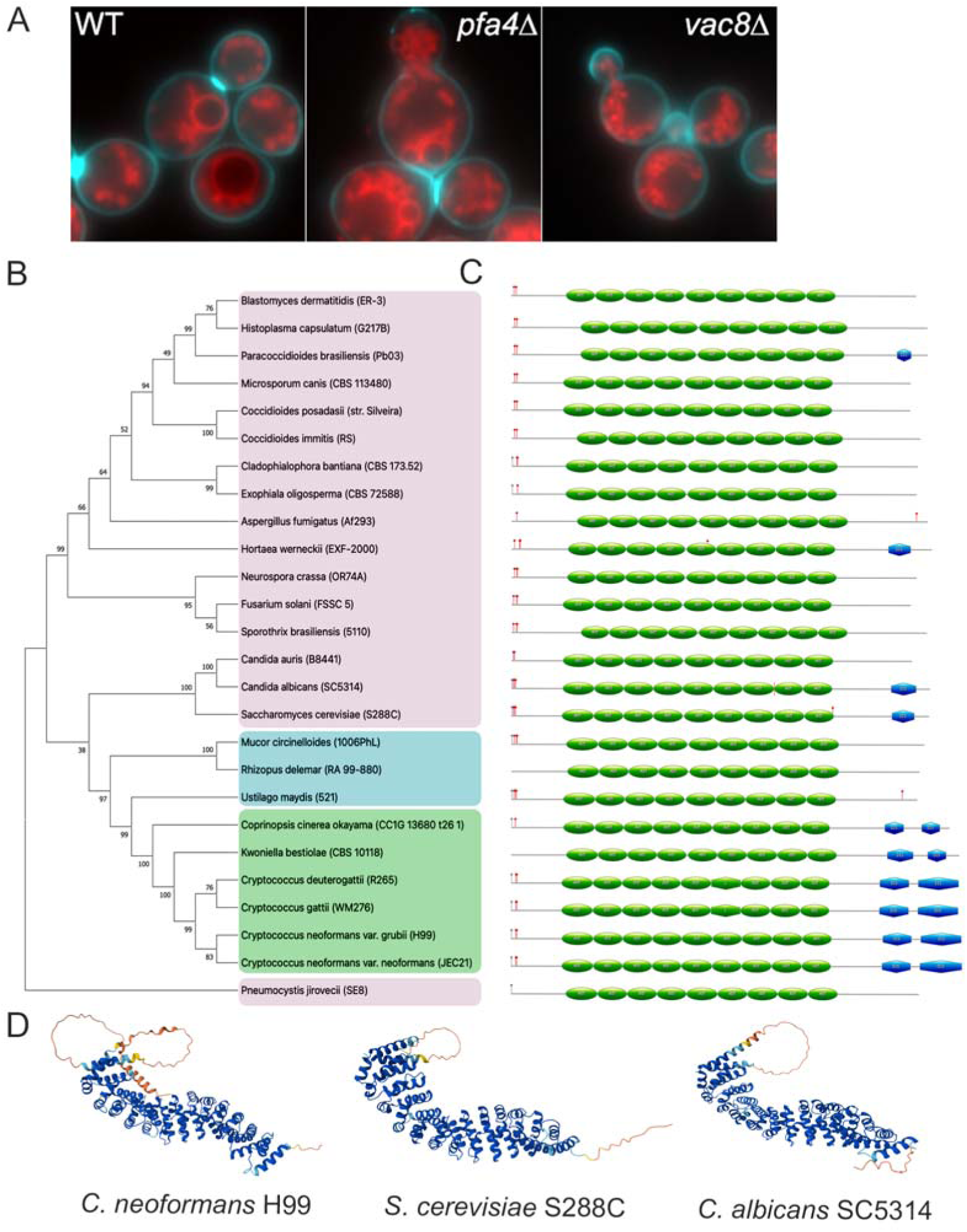
CNAG_00354 is a true homolog of ScVAC8. (**A**) Vacuole localization and morphology in wild- type (*23*), strains lacking palmitoylation of Vac8 (*pfa4Δ*), or lacking Vac8 itself (*vac8Δ*). Vacuoles were stained with FM4-64 (*59*) and the cell outline with calcofluor white (blue). (**B**) Phylogenetic tree depicting the Vac8 amino acid sequence alignments in many clinically relevant fungi. Node support is represented as a percentage of 1000 bootstrapping iterations. Magenta shading: ascomycetes, blue shading: Mucorales, green shading: basidiomycetes. (**C**) Graphic representation of protein structure based on InterPro analysis of amino acid sequence shown to scale, relative to other Vac8 proteins. 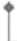 indicates predicted myristylation site, 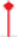 indicates predicted palmitoylation site, green ovals represent armadillo repeat regions, blue hexagons represent predicted disorder regions. (**D**) Alphafold predictions of *C. neoformans*, *S. cerevisiae*, and *C. albicans* Vac8.

**Figure 2:**
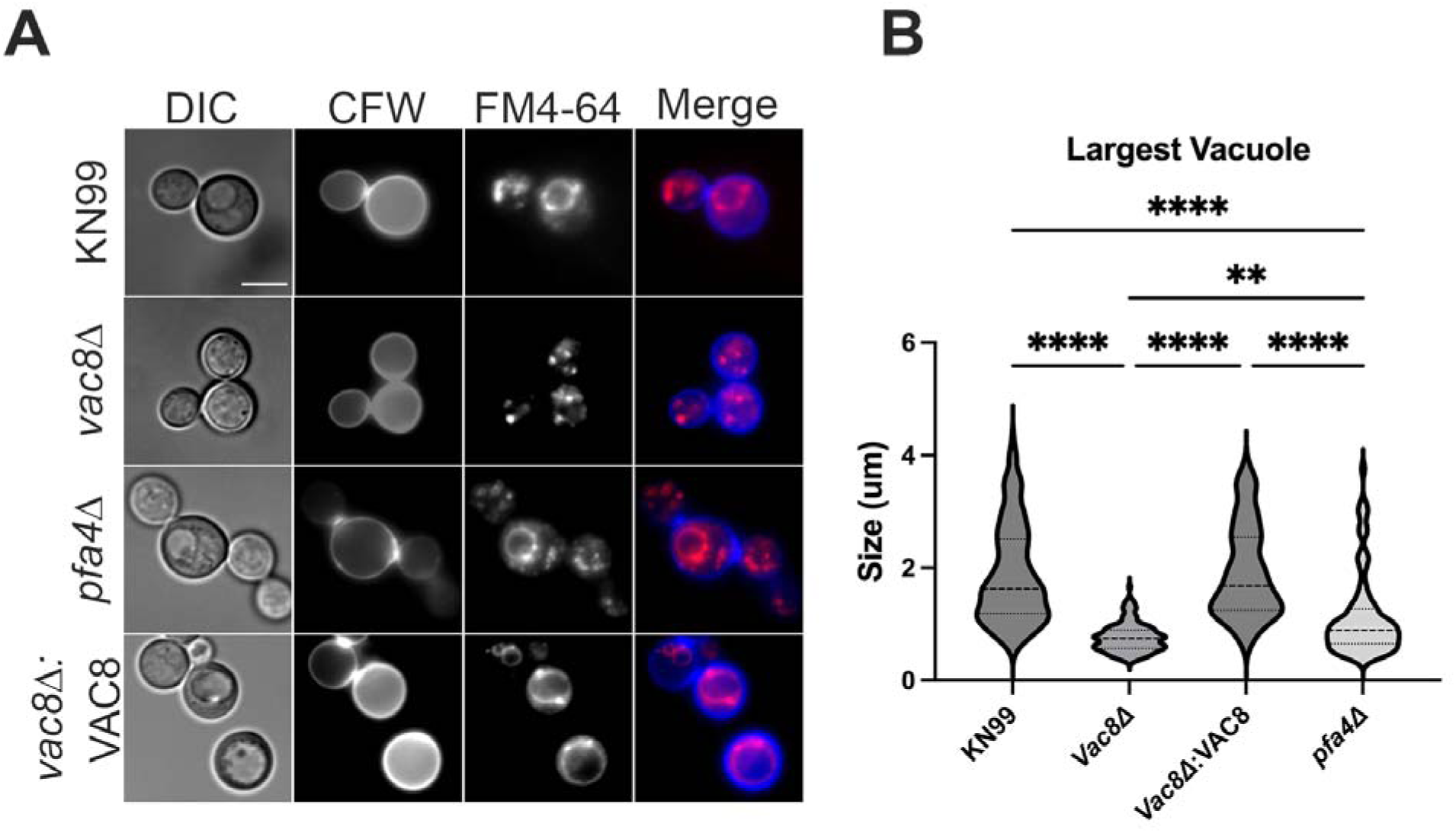
Vac8 regulates vacuole fusion. (**A**) Cells were cultured in YPD at 37°C supplemented with 1M sorbitol for 4 hours before being stained with calcofluor white and FM4-64 to visualize cell wall and vacuole, respectively. Scale bar: 5 μm. (**B**) Quantification of the largest vacuole in each cell by diameter (n: 84-246). *vac8Δ* strains and *pfa4Δ* all show defects in ability to form a large vacuole under thermal and osmotic stress. Significance was assessed by 2-way ANOVA with multiple comparisons, ** = p=0.001, **** p<0.0001.

### CNAG_00354 is a true ortholog of Saccharomyces cerevisiae *VAC8*

Given the similar but not identical phenotypes relative to *S. cerevisiae*, we sought to determine if CNAG_00354 is a true orthologue of *ScVAC8* by performing reciprocal BLAST-P searches. We found a 1:1 relationship if using CNAG_00354 to search the *S. cerevisiae* genome or *ScVAC8* to search the *C. neoformans* genome, confirming that CNAG_00354 (from now on named *VAC8*) is a true ortholog.

Next, we performed a phylogenetic analysis to determine sequence similarity and how widespread this protein is in the fungal kingdom (**Figure 1B**). The basidiomycetes, ascomycetes, and Mucorales species all cluster together within their respective groups; while the basidiomycetes and Mucorales appear to be more closely related than ascomycetes are to either clade. This clustering indicates *VAC8* is well conserved in the fungal kingdom but there are substantial differences from Ascomycota, including *S. cerevisiae* which most of our fungal knowledge is based on. This suggests there may be both similar as well as unique functions of *VAC8* in *C. neoformans* and other basidiomycetes or Mucorales compared to ascomycetes.

In *S. cerevisiae*, Vac8 is tethered to the vacuole membrane by an N-terminal glycine that is myristoylated and three cysteines near the N-terminus that are palmitoylated, allowing the protein to anchor the fungal vacuole.(*29*) Alphafold prediction indicates highly conserved structure between ScVac8 and Vac8 including an N-terminal tail with glycine and cysteine residues (**Figure 1D**). Using the palmitoylation prediction software GPS Palm (*24*) and myristylation software (*25*), we investigated the likelihood of the cysteines in Vac8 being palmitoylated and the N-terminal glycine being myristoylated. In ScVac8 the cysteine residues at position 4, 5, and 7 are known to be palmitoylated (Error! Reference source not found.). Vac8 has cysteine residues in positions 8 and 9 that are also predicted to be palmitoylated (Error! Reference source not found.).(*24*) In congruence with the phylogenetic tree, *Cryptococcus gattii* has cysteine residues in positions 8 and 9 with predicted palmitoylation values 0.9714 and 0.9667 while *Ustilago maydis* has cysteine residues in positions 5, 7 and 8 with predicted palmitoylation values 0.9677, 0.9396, and 0.9300 respectively, while the glycine is also myristoylated (**Figure 1B,C**).(*24*) Taken together, these data indicates that the overall protein domains and modifications of *VAC8* are conserved across the fungal kingdom and *C. neoformans VAC8* is a true ortholog of *ScVAC8*.

Vac8 is an ARM repeat protein, comprised of the characteristic repetitive amino acid sequence of approximately 42 residues. These residues are often tandemly repeated where each repeat gives rise to a pair of alpha helices which form a hairpin structure. Crystal structures have revealed ScVac8 has 11 complete ARM repeats, and an incomplete 12th ARM repeat which form alpha helices that fold to form a superhelix allowing for Vac8 to interact with a variety of binding partners.(*30*) This diversity in Vac8-protein interactions is what allows it to carry out diverse cellular functions. To this end we explored the *C. neoformans* genome for orthologues of known ScVac8 binding partners. Many of the main known binding partners of ScVac8, including Nvj1, Vac17, Atg13, and Tco89 do not have homologues in *C. neoformans*, while Tao3 (transcriptional activator of *OCH3*) involved in apical bud growth and morphogenesis, is one of the few proteins with a potential homologue (Error! Reference source not found.).

### Deletion of *VAC8* affects vacuolar fusion

Since deletion of *PFA4* and *VAC8* results in fragmented vacuoles under nonstress conditions (YPD and 30°C; **Figure 1A**), we hypothesize that *VAC8* might be playing a direct role in vacuolar fusion. To test this, we visualized vacuoles under conditions known to induce large vacuole formation in WT *C. neoformans* (YPD+1M Sorbitol and 37°C).(*28*) We saw WT *C. neoformans* KN99 have large, readily visible vacuoles in both FM4-64 and DIC imaging, while *pfa4Δ* has considerably smaller vacuoles with relatively few cells generating a vacuole large enough to be seen in DIC. (Error! Reference source not found.) *vac8Δ* also showed a marked inability to generate large vacuoles in response to high temperature and osmotic stress, with no obvious vacuoles being observed through DIC. (Error! Reference source not found.) Average size of vacuoles was compared between each strain by measuring the diameter of the largest vacuole as observed by FM4-64 staining and DIC. Loss of *VAC8* was sufficient to inhibit vacuole fusion under set conditions. (Error! Reference source not found.)

### Loss of *VAC8* is associated with growth defects in caffeine media, cell membrane and cell wall stresses

Given the importance of normal vacuole function in a variety of cellular processes, *vac8Δ* was subjected to a series of cellular stressors. All strains tested grew equally well on both YPD agar at 30°C. In *S. cerevisiae*, when Vac8 was put under the Vac17 promoter resulting in 10-fold lower *VAC8* expression, there were increased levels of β-1,3-glucans and β-1,6-glucans, two of the major fungal cell wall components. (*31*). The *pfa4Δ* mutant, which lacks the palmitoyl acyl transferase for Vac8, displayed growth defects in high temperature (37°C and 37°C with 5% CO_2_), high salt (1M NaCl), and caffeine (0.75 mg/mL caffeine) as previously demonstrated (**Figure 3**). *vac8Δ* also showed significantly reduced growth at 0.75 mg/ml caffeine suggesting an impaired response to general cell stress. (**Figure 3**) Interestingly, growth on 2 mM H_2_O_2_ YPD plates is comparable to that of WT, but on 4 mM H_2_O_2_ *vac8Δ* exhibits markedly increased growth, while *pfa4Δ* and KN99 growth is limited (**Figure 3**).

**Figure 3:**
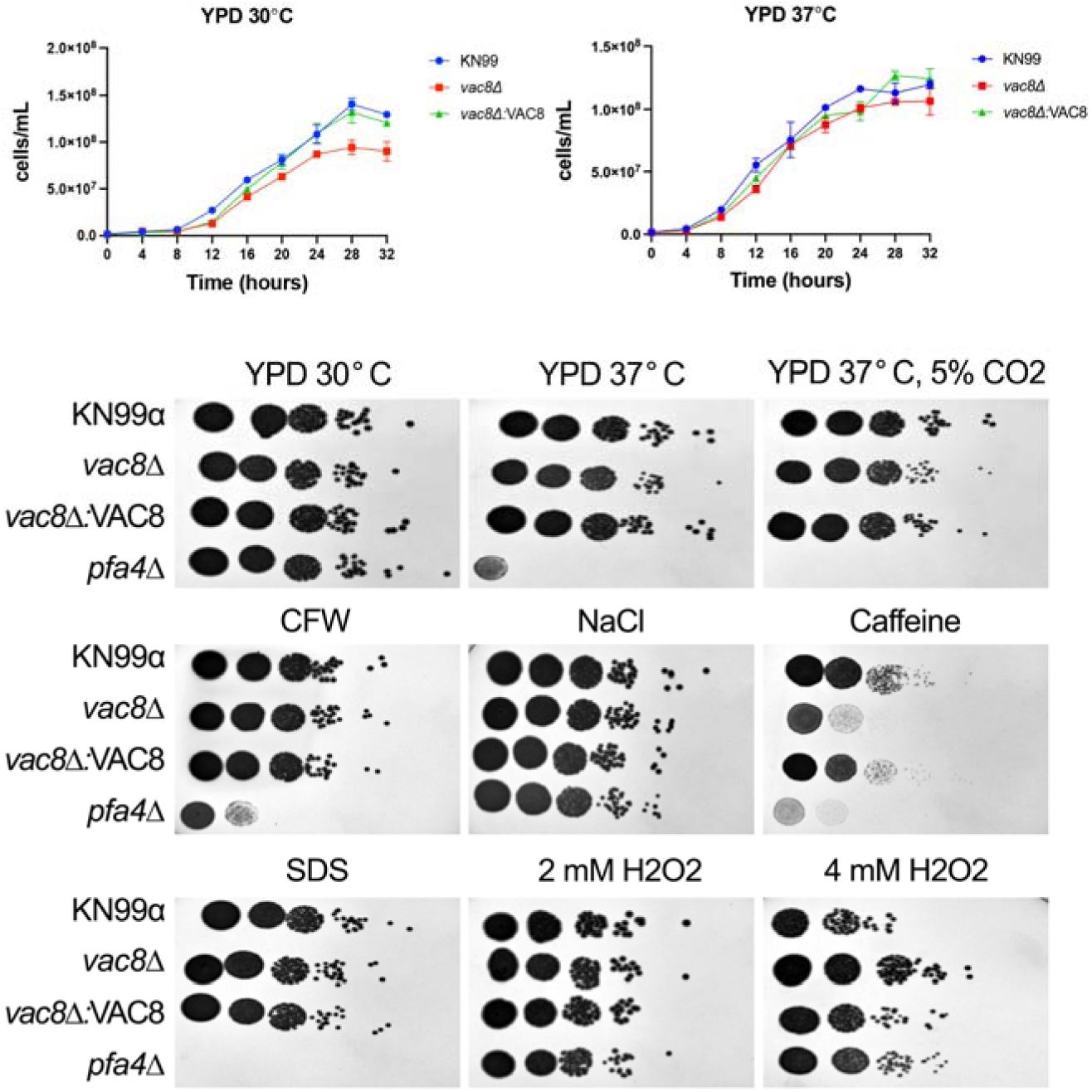
*vac8Δ* displays increased sensitivity to cell wall stressors and caffeine but increased resistance to oxidative stress. Growth curves showing cell counts of KN99, *vac8Δ*, and *vac8Δ*:VAC8 grown in YPD at 30 or 37° C for 32 hours. Serial dilutions (10^7^ to 10^3^) of cells were spotted on media containing the following stressors: .03% SDS, 1M sorbitol, 2 or 4 mM H_2_O_2_, 0.5 mg/mL CFW, 1M NaCl, or .75 mg/mL caffeine. Images shown are representative of at least two independent replicates.

### Main virulence factors are unaffected in *vac8Δ* mutants

Previous studies in *C. neoformans* and *C. albicans* have shown that normal vacuole function is needed for full virulence.(*10, 11, 32*) Similarly, in other human fungal pathogens like *Candida glabrata* and *Histoplasma capsulatum*, normal vacuolar function has been implicated in the ability to cause disease through the vacuole’s role in autophagy and iron homeostasis.(*13, 14*) To explore whether deletion of *VAC8* causes a defect in any of the main *C. neoformans* virulence factors we tested growth at high temperature, capsule production, melanin production, and urease activity in WT and *vac8Δ*.(*33*) *vac8Δ* displays a slight decrease in growth on YPD agar at 37° C and 37° C with 5% CO_2_ suggesting it may have reduced fitness in an infection model (**Figure 3**). Additionally, *vac8Δ* shows slower growth rates at 30° C relative to KN99 (**Figure 3**).

Further we explored the effect of *vac8Δ* on induction of the cryptococcal capsule as this is a critical virulence factor involved in resisting the effects of the host innate immune system.(*34*) While some *C. neoformans* vacuolar mutants like *vph1Δ* do not produce capsule compared to WT, *vac8Δ* displays a modest but significant reduction in capsule but still produced substantial levels of capsule (**Figure 4A-C**). Laccase activity was assessed by melanin production in the presence of L-DOPA media and all mutants displayed normal melanization (**Figure 4D**). Urease activity, determined by a phenol red-indicated pH change in the presence of urea, appeared reduced after 7 days in *vac8Δ* compared to KN99 and *vac8Δ*:*VAC8*. (**Figure 4D**).

**Figure 4:**
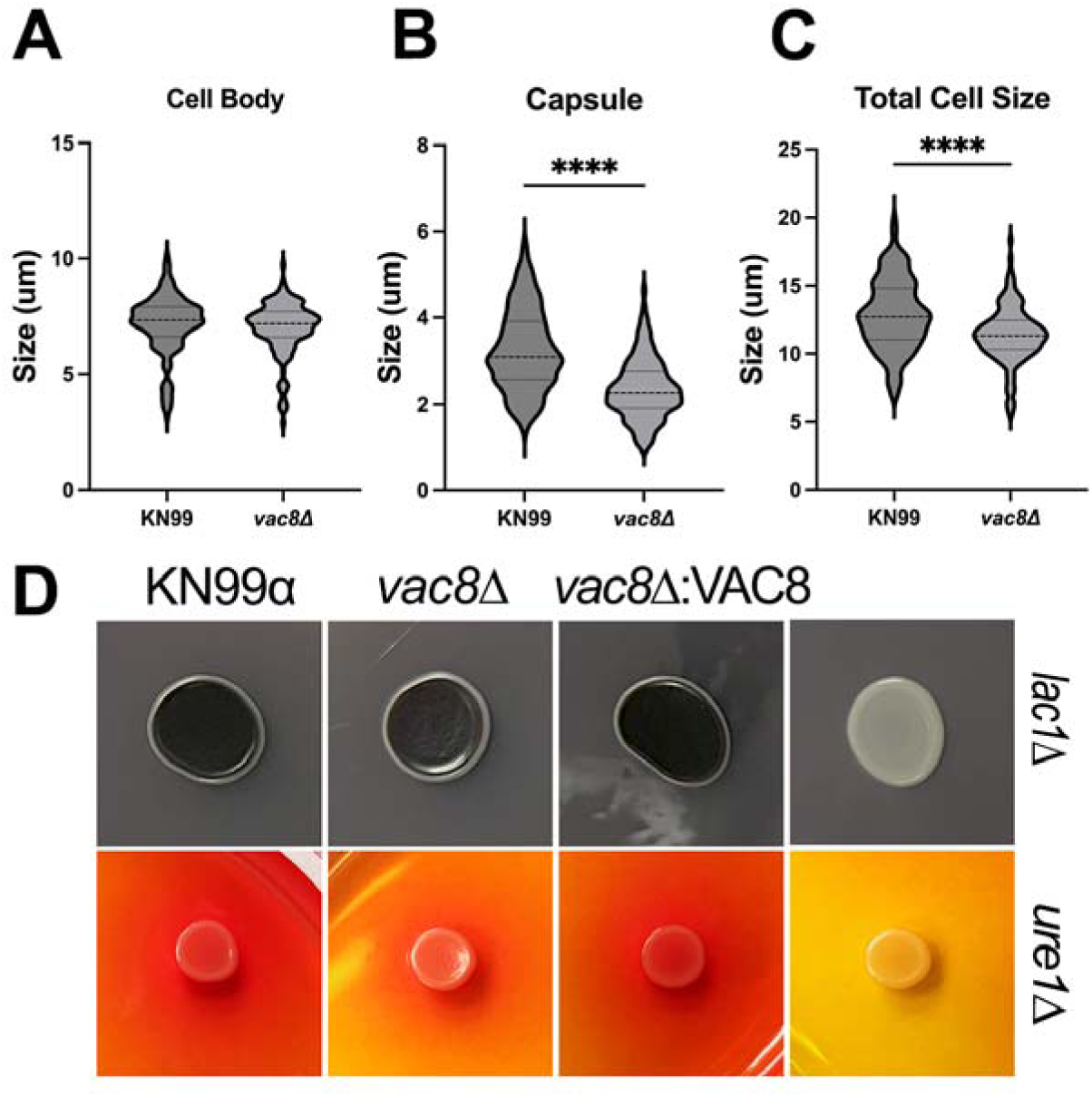
*vac8Δ* demonstrates a slight reduction in urease activity and reduced ability to form capsule which results in a reduced overall cell size. (**A-C**) Quantification of cell body (**A**), capsule (**B**), and total cell size (**C**) in capsule inducing media (DMEM, 5% CO2, 37C) after 48 hours. For each measurement, at least 120 cells were analyzed. (**D**) Melanin production on L-DOPA plates and 168 hours and Urease production on Christensen’s Urea agar plates at 168 hours.

### *vac8Δ* mutants show aberrant budding and impairment in titanization

Morphological transitions are a hallmark of fungal adaptations to host environments and play an important role in disease. In *C. neoformans*, formation of the polysaccharide capsule and titan cell formation are regulated by the cell cycle. (*35–38*) The cryptococcal capsule is one of the best characterized virulence factors of *C. neoformans*, and it is critical for infection as acapsular mutants display severely attenuated virulence.(*34*) Under capsule inducing conditions (37°C + 5% CO_2_, DMEM, and 48 hours) *vac8Δ* displayed increased levels of aberrant budding morphology with incomplete cytokinesis including elongated cells and mother cells with multiple daughter cells branching from them (**Figure 5**). As incomplete cytokinesis is tied to abnormalities in the cell cycle, we sought to determine if the number of cells with abnormal budding morphology increased over time. To explore this, we examined the bud morphology of cells grown in DMEM at 37° C + 5% CO_2_ every 24 hours for 96 hours. KN99 displays little to no cells with more than one bud at all time points examined (**Figure 5**). Interestingly, the percentage of cells with aberrant budding morphology did not increase over time in *vac8Δ*, yet *vac8Δ* had significantly more budding defects compared to wildtype strains at each time point examined (**Figure 5**).

**Figure 5:**
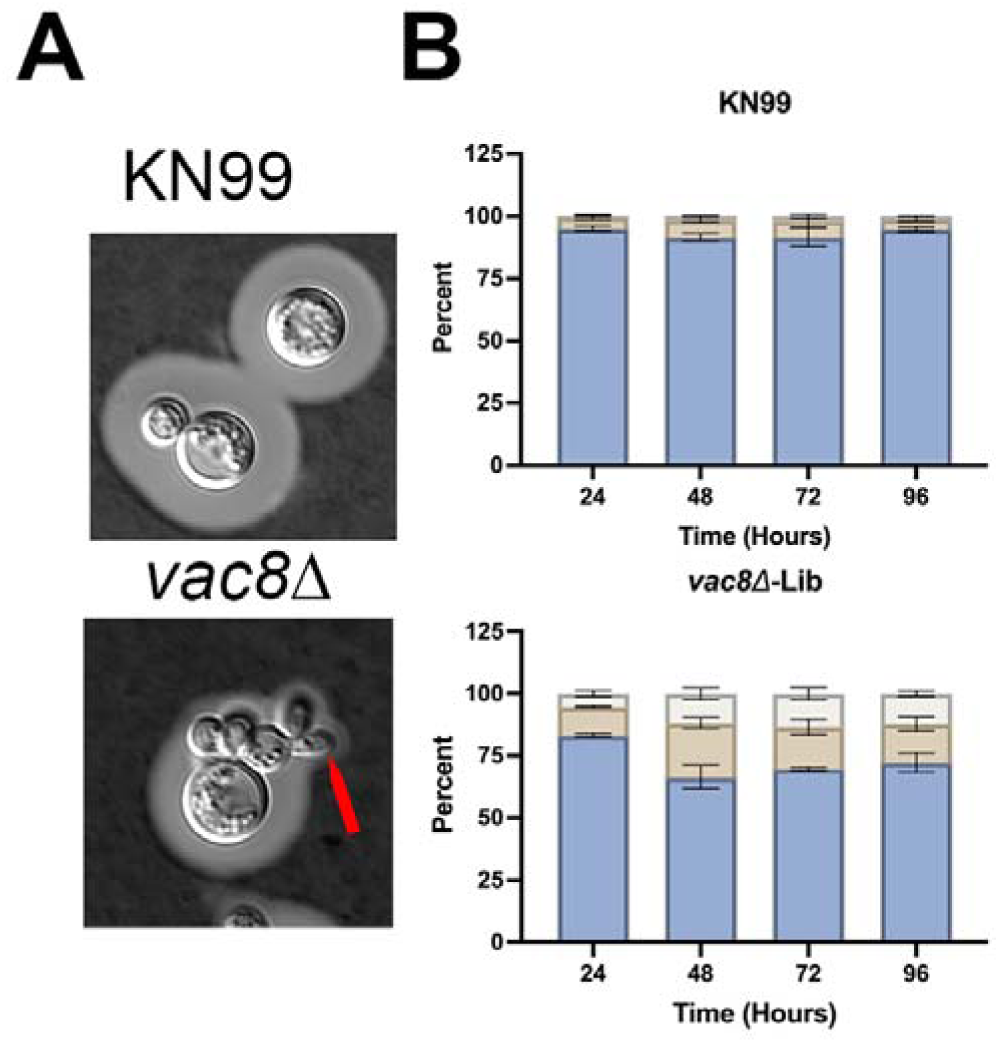
*vac8Δ* mutants show aberrant budding morphology. (A) DIC Images of wildtype and *vac8Δ* strains after 48h in capsule inducing conditions stained with India ink. Red arrow indicates elongated bud morphology. **(B**) Quantification of budding morphology over time as determined by the total number of buds stemming from a single mother cell. Data represents percentages and SEM of 2 independent replicates. 150-700 cells were measured per timepoint per condition.

Titan cells are a unique morphological state that can be induced *in vivo* by mammalian lung environment and internal *Galleria mellonella* environment.(*37–39*) These cells are characterized by their large size. Normal *C. neoformans* yeast cells have a cell body 5-7 μm while cell bodies of titan cells can reach up to 100 μm; an arbitrary cut off of either 10 or 15 μm is commonly used for classification.(*40*) Additionally, these cells display increased ploidy, often tetraploid up to 128-ploid compared to the haploid yeast, a large vacuole that encompasses the majority of intracellular space, a thickened cell wall (2-3 μm vs 150-200 nm), and a denser more cross-linked capsule.(*40*) These cells play a critical role during infection by contributing to fungal survival, promote dissemination to the CNS, and are involved in latency and resistance to antifungal treatment.(*40, 41*) Given that large vacuole formation is observed in titan cells and *vac8Δ* shows fragmented vacuoles, we sought to determine if *vac8Δ* can generate titan cells. Recently, multiple groups have identified ways to create *bona fide* titan cells in vitro using low nutrient, serum supplemented medium at neutral pH with CO_2_.(*37, 39*) We utilized the Dambuza *et al.* method where cells are cultured overnight in YNB, washed, and diluted to 1000 cells/mL in 10% heat inactivated FBS and incubated at 5% CO_2_, 37° C for 48 hours. Using a 10 μm threshold for cell body, we saw results consistent with previous studies where ∼10% of KN99 cells titanized, significantly less than *usv101Δ* at 42.7% which readily titanizes, (**Figure 6A,B**). *vac8Δ* showed greatly reduced ability to form titan cells at 1.1%, titanization. Despite the inability to form titan cells, *vac8Δ* cells are able to produce a robust capsule albeit smaller than KN99 and complement strains (**Figure 6C**). This led to WT strains and *usv101Δ* having significantly larger cells overall, approaching 40 μms, while very few *vac8Δ* cells reached 20 μms total cell size (cell body and capsule) (**Figure 6D**).

**Figure 6:**
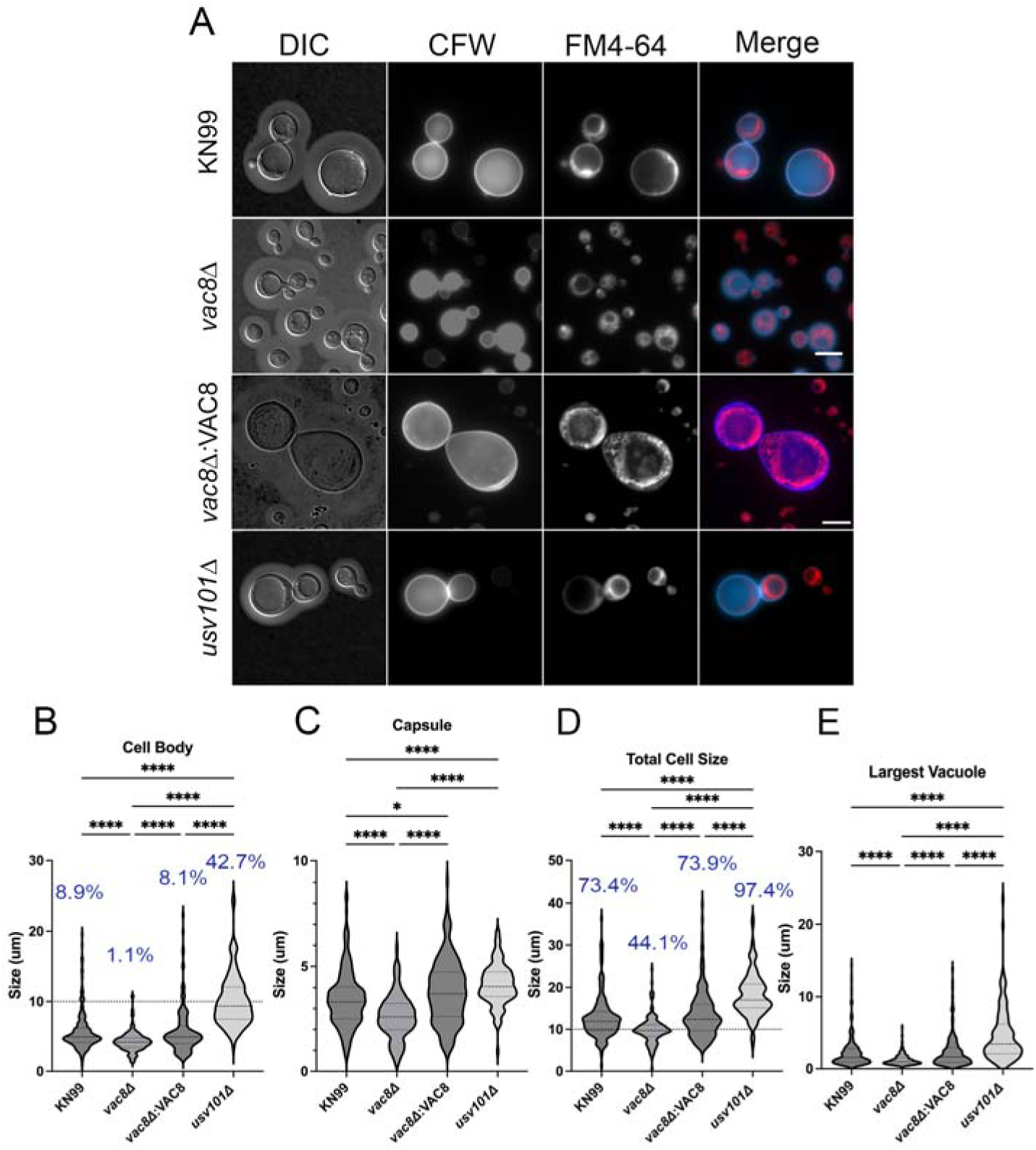
Loss of *VAC8* reduces the ability to form titan cells and induces aberrant budding morphology in titan cell inducing conditions. (**A**) Images of cells grow in titan inducing conditions (10% FBS, 5% CO_2_, 37° C) for 48 hours. Cells were stained with calcofluor white (CFW) and FM4-64 to visualize cell wall and vacuole structures, respectively. (**B – E**) Quantification of cell body (**B**), capsule (**C**), total cell size (**D**) and the largest vacuole in the mother cell. Scale bar is 10 μms. Dashed line represents 10 μm titan cell threshold. Blue percentages represent the percentage of cells > 10 μm. At least, 115 cells were analyzed for each strain across 2 independent experiments. Significance was determined by ordinary one-way ANOVA with multiple comparisons. *, P < 0.05; ****, P < 0.0001.

Internally, titan cells produce a single large vacuole that may be involved in extracellular vesicle production.(*7*) It is unclear if the vacuole of titan cells behaves similarly or differs from that of typical cells. Here, we observed KN99 and *usv101Δ* typically display a single central vacuole that takes up the vast majority of the cell and can easily be seen in DIC imaging or with the lipophilic dye FM4-64 (**Figure 6A**). In contrast, *vac8Δ* cells often show multiple smaller vacuoles that make up less of the internal environment, even in *vac8Δ* cells that would be considered titan cells (**Figure 6A**). Given this, the ability to generate titan cells may be dependent on the ability to form a single large vacuole. (*9, 11*)

### Loss of *VAC8* casuses decreased susceptibility to fluconazole

Due to the close evolutionary relationship between humans and yeast, there are few antifungal targets within yeast, which has resulted in the very limited number of antifungal drugs.(*2*) Most antifungal classes target the biosynthesis of ergosterol or the ergosterol molecule itself, the main sterol in fungal membranes.(*2, 42*) While a newer class of antifungal, the echinocandins, target biosynthesis of (1,3)-β-glucan, a key component of the fungal cell wall, it has limited applicability and in fact is ineffective against *Cryptococcus*.(*2*) Therefore, there is a clear need to find novel drug targets, specifically in *C. neoformans*. Vacuoles play an important role in the detoxification and sequestration of compounds in yeast. Recently, they have also been implicated in fluconazole resistance in *C. albicans* and *S. cerevisiae* through the ability of ABC transporters to import fluconazole into the vacuole.(*43–45*) Additionally, as fluconazole empties vacuolar membranes of ergosterol, or amphotericin B binds to ergosterol there, it inhibits V-ATPase complex formation, preventing the vacuole from acidifying.(*46, 47*)

To this end, we examined if *vac8Δ* and other mutants (*pfa4Δ* and *flc1Δ*), with their abnormal vacuolar morphology, display increased susceptibility to two common antifungals: fluconazole (azole class) and amphotericin B (polyene). First, cells were incubated under optimal conditions (YPD and 30° C) to determine if aberrant vacuole morphology affects growth under these conditions. (**Figure 7**) While no strain showed impaired growth on YPD, growth at high temperature on RPMI plates with impaired in *vac8Δ*, *pfa4Δ*, and *flc1Δ*. Interestingly, the addition of CO_2_ slightly recovered the growth of *pfa4Δ* while not markedly changing the relative growth of *vac8Δ* or *pfa4Δ*. (**Figure 7**)

**Figure 7:**
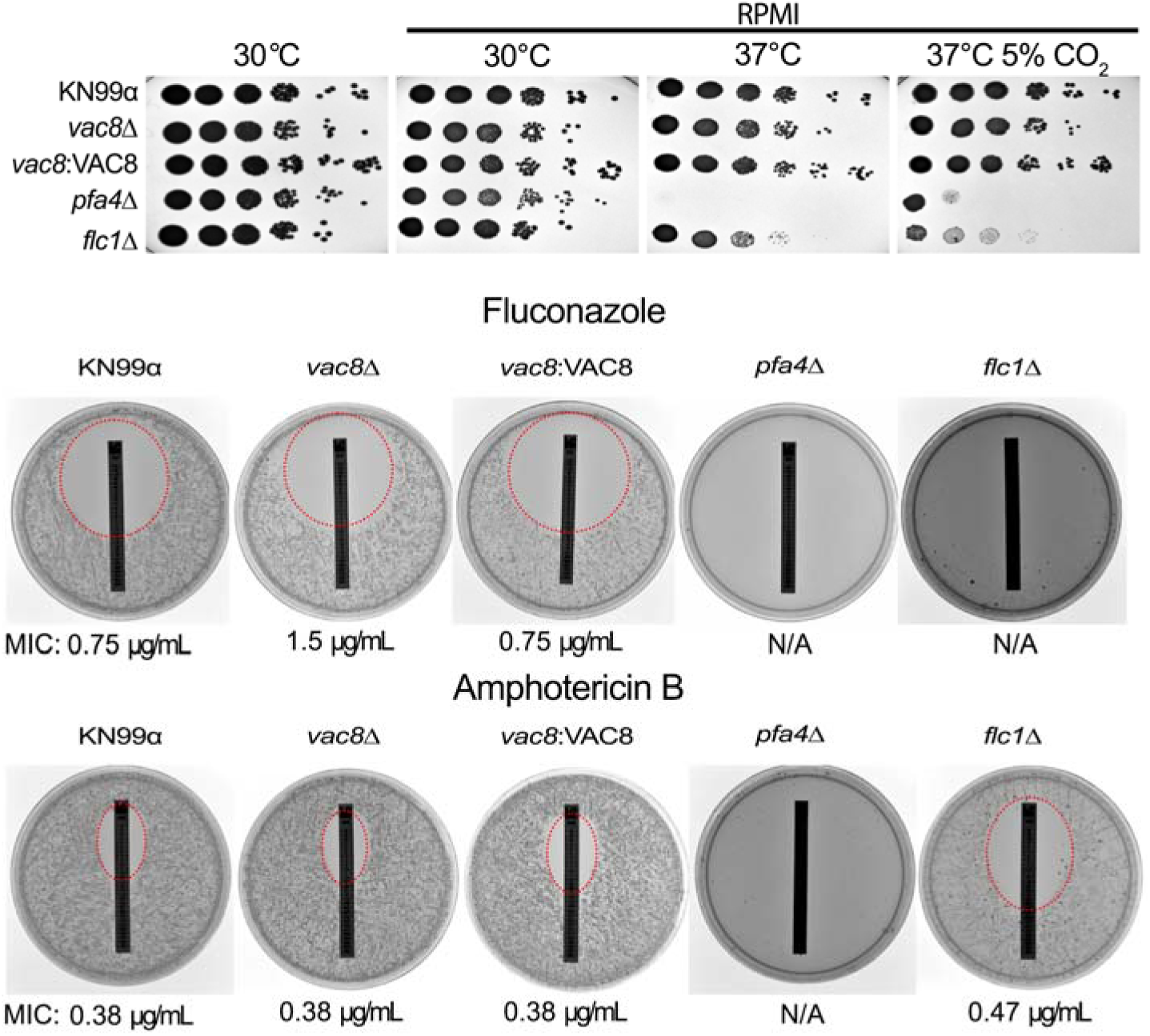
*vac8Δ* strains show a slight increase in resistance to fluconazole but no changes to polyene susceptibility despite a slight sensitivity to high temperature and CO_2_ growth. The indicated strains were grown plated in serial dilution on YPD, or RPMI plates and incubated at 30° C, 37° C, or 37° C with 5% CO_2_ for 72 hours before being imaged. FLC or AmpB Epsilometer test strips were applied to RPMI 1640 agar plates with 2 × 10^5^ cells of the indicated strain. The zone of inhibition is highlighted with a red dashed circle where applicable with the MIC listed below the image. The MICs of KN99 and *vac8Δ* mutants were all 0.19 μg/mL. *pfa4Δ* shows complete inhibition of growth in the presence for both FLC and AmpB, while *flc1Δ* is completely inhibited by FLC and has an MIC of 0.47 μg/mL for AmpB. These plates were incubated for 72 h at 37° C and 5% CO_2_.

To examine susceptibility to antifungals cells were grown overnight in YPD then washed twice with PBS and 2x10^5^ cells were spread across RPMI agar plates and allowed to dry prior to application of an E-test strip. After 72 hours, minimum inhibitory concentrations (MICs) were determined by examining where the edge of the zone of inhibition borders the test strip. Interestingly, the *vac8Δ* strain showed a modest decrease in susceptibility to fluconazole, with MICs of 1.5 μg/mL compared to .75 μg/mL for KN99 (**Figure 7**). The MIC for AmpB remained consistent at 0.38 μg/mL (**Figure 7**).

### Infection with *vac8Δ* does not alter disease outcome in a murine model

Innate lung immune cells, in particular alveolar macrophages represent the first line of defense against cryptococcal infections, and the complex fungal-macrophage interaction will ultimately help determine the outcome of the infection. (*48*) The *vac8Δ* mutant has previously been indicated as high uptake in both murine and human bone marrow derived macrophages. (*49*) To determine if *vac8Δ* cells display different interactions with alveolar macrophages *in vitro*, cells were incubated with THP-1 macrophages for one hour and phagocytic index was determined as previously reported. (*19*) There was no difference in uptake of KN99, *vac8Δ*, and *vac8Δ*:*VAC8* cells by THP-1 cells. (**Figure 8A**). To further explore to role of *vac8Δ* in *C. neoformans* pathology, 5-6 week old A/J mice were infected with 5x10^5^ cells via intranasal inoculation. Mice infected with *vac8Δ* started to succumb to infection prior to KN99 and *vac8Δ*:*VAC8* infected mice but mean survival was not affected. (**Figure 8B**) Given the difference in responses to cellular stressors, fungal organ burden was determined during at 3-, 7-, and 14-dpi. At 3- and 7- dpi no strain had disseminated to the spleen. (**Figure 8C-D**) At 3-dpi both KN99 and *vac8Δ*:*VAC8* infected mice showed low levels of dissemination to the brain, while only the *vac8Δ* infected mice showed low levels of dissemination at 7-dpi. (**Figure 8C-D**) Fungal organ burden in the brain, spleen, and lung was similar amongst mice infected with each strain at all time points. (**Figure 8C-F**)

**Figure 8:**
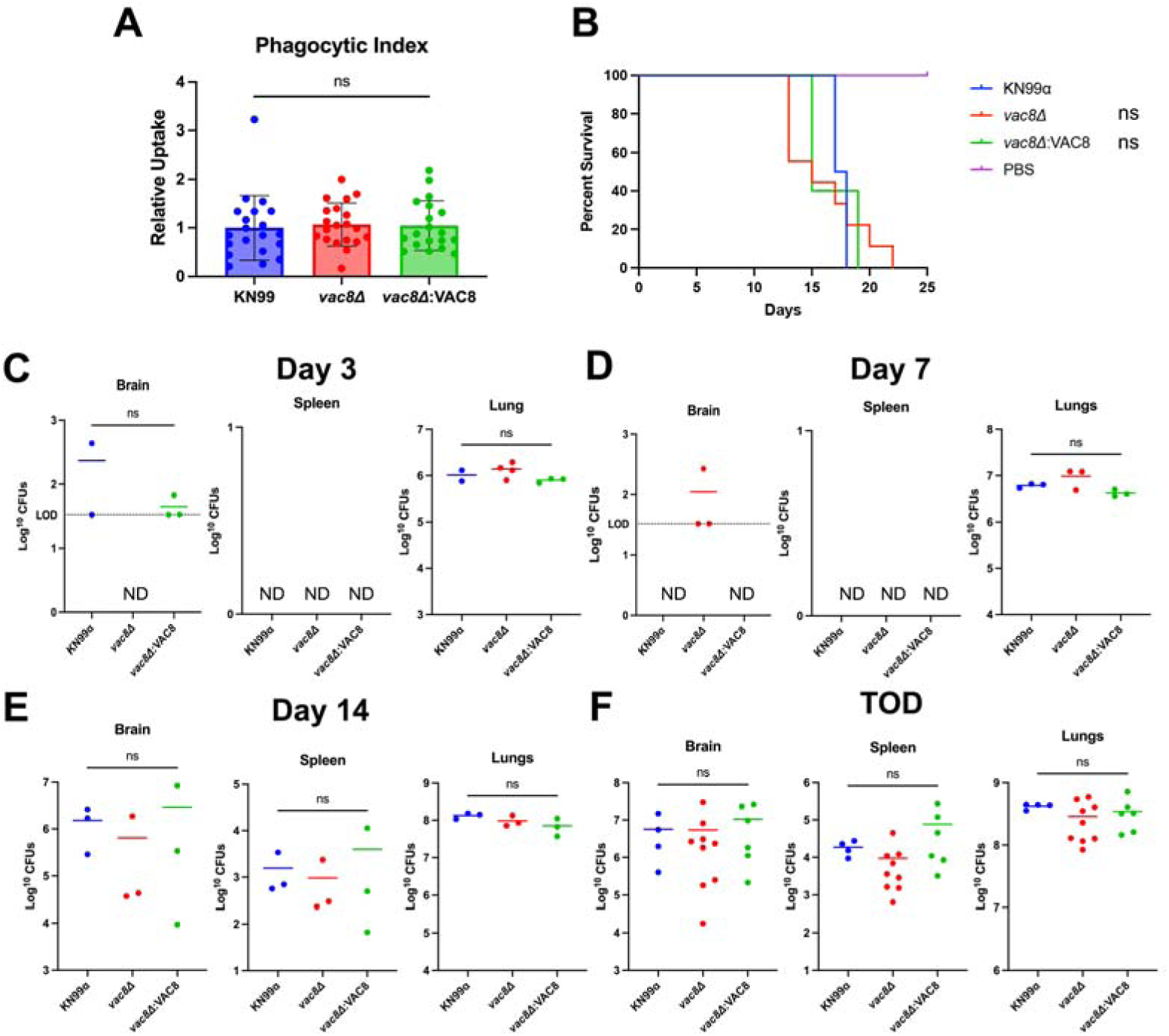
*C neoformans vac8Δ*-infected mice succumb to infection at similar rates to KN99- infected mice. (**A**) Phagocytic index of THP-1 macrophages infected with the indicated fungal strains at an MOI or 1. Significance was determined using an ordinary one-way ANOVA with multiple comparisons. (**B**) *In vivo* survival of 5-6 week old female A/J mice infected with KN99, *vac8Δ*, *vac8Δ*:VAC8 or DPBS (Mock) and monitored for 25 days when the experiment was terminated. 5-10 animals were used per group. Significance was determined using Mantel-Cox test. (**C-F**) Quantified organ burdens of KN99, *vac8Δ*, and *vac8Δ*:VAC8-infected mice at 3-, 7-, and 14-days post infection, and at time of death (TOD). Organs were harvested, homogenized, and dilutions were plated on YPD plates. Plates were incubated for 48 h and CFUs were counted. Each data point represents a mouse, and black bars represent the mean. Significance was determined using an ordinary one-way ANOVA with multiple comparisons.

### Infection with *vac8Δ* alters the lung environment despite not impacting disease progression

*Cryptococcus* initially colonizes the lung environment, yet the fatal pathology is associated with dissemination to the central nervous system and meningoencephalitis. (*50, 51*) Therefore the lung environment is critical in controlling the overall course of infection. To qualitatively examine any differences at the tissue and cellular level, lungs of mock, KN99, *vac8Δ*, and *vac8Δ*:*VAC8*-infected mice were harvested and stained with hematoxylin and eosin (H&E). By 7-dpi the lungs of *vac8Δ* infected mice showed increased pockets of inflammation that was absent from lungs of mice infected with KN99, *vac8Δ*:*VAC8*, or mock. (**Figure 9A**) This trend continued at 14-dpi with all infected lungs displaying some level of tissue damage. (**Figure 9A**) Despite not seeing differences in organ burden by CFUs, *vac8Δ* infected lungs showed pockets of inflammation near blood vessels and large nodules in the lung where all healthy tissue had been destroyed leaving large pockets filled with yeast. (**Figure 9B**) With similar levels of fungal burden the lungs of KN99 and *vac8Δ*:*VAC8* infected mice show minimal levels of inflammation, characteristic of cryptococcosis. (**Figure 9B**) Interestingly, the yeast observed in KN99 and *vac8Δ*:*VAC8* infected lungs are mostly single celled and very few appear to have a bud, while *vac8Δ* cells in the lungs display similar aberrant morphologies seen in vitro. (**Figure 5**, **Figure 9B** Red inset)

**Figure 9:**
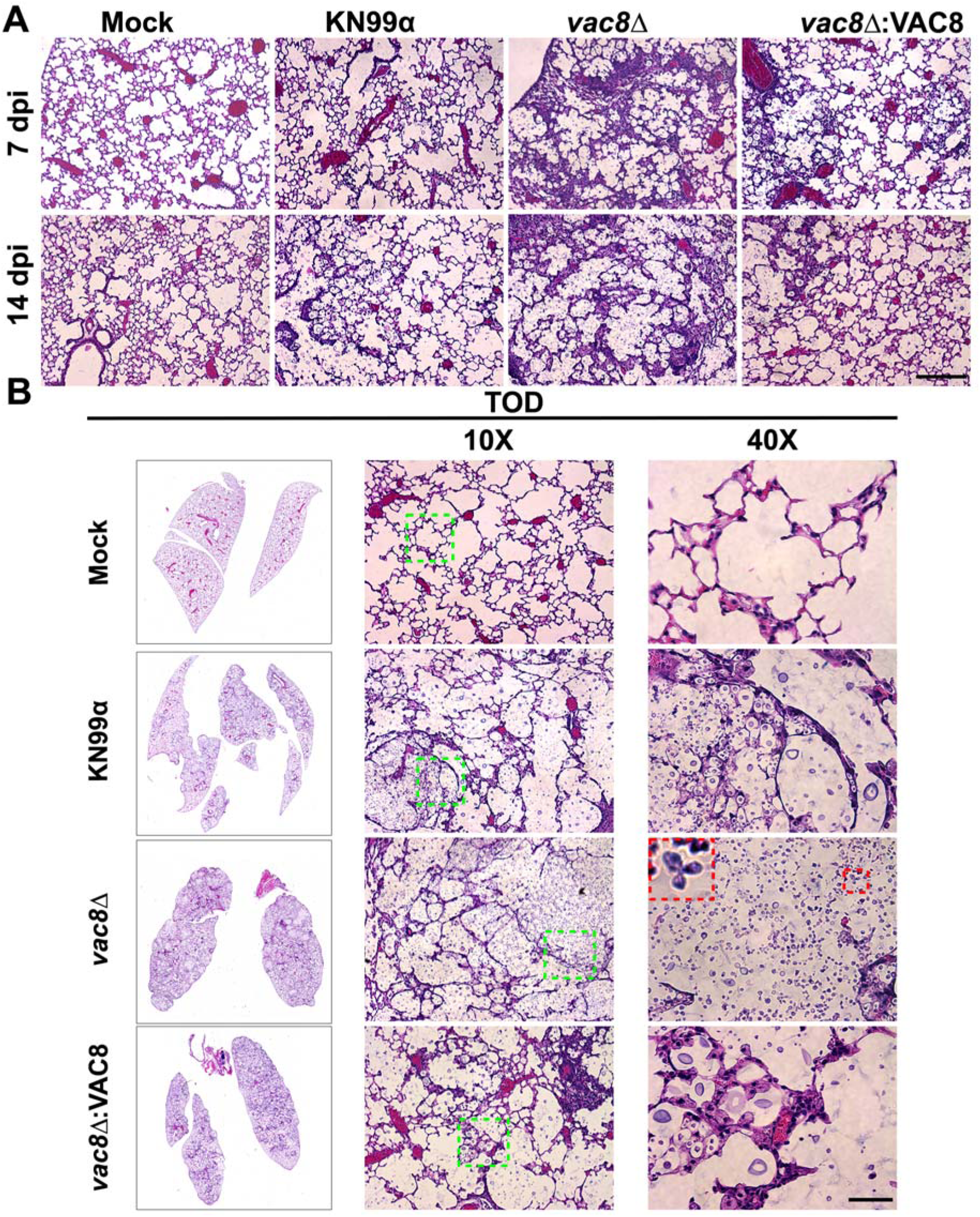
Infection with *C. neoformans vac8Δ* results in small differences in histopathology. (**A**) Dissected lungs of female A/J mice infected with 5 x 10^4^ cells of the KN99 wildtype strain, *vac8Δ*, or *vac8Δ*:VAC8 strain or DPBS infected (Mock) were harvested at predetermined endpoints of 7 and 14 dpi. (**B**) Gross organ examination revealed significant changes in inflammation and tissue damage at time of death (TOD). H&E staining was used to visualize microscopic lung pathology. Green boxes on 10X images are inset as 40X. Red box highlights abnormal fungal morphology observed in the lungs of mice infected with the *vac8Δ* strain. The 40X scale bar is 50 μm.

## DISCUSSION

Here, we report an initial characterization of vacuolar protein 8 of *C. neoformans* and its role in vacuolar and cellular morphologies in response to cell stress. First, we confirmed that CNAG_00354 is a true homologue of ScVac8 with conserved myristylation and palmitoylation sites at the N-terminus to allow for association onto the vacuolar membrane, and vacuolar defects when deleted. (**Figure 1**) Additionally, Interpro and Alphafold predictions reveal a protein structure very similar to the ScVac8 crystal structure, with a central superhelix comprised of 11 ARM repeats serving as a protein binding platform, a lipid modified tail that attaches it to membranes, and a disordered region near the C- terminus. Despite a far evolutionary distance between ascomycetes, basidiomycetes, and Mucorales species, Vac8 has a well conserved protein structure that explains the overlapping and distinct roles it plays over evolution.

In *S. cerevisiae* Vac8 is well characterized and has multiple binding partners which facilitate its involvement in key cellular processes like piecemeal autophagy of the nucleus via the nuclear-vacuole junction, autophagy via Atg13, and vacuolar inheritance via Vac17. Given the evolutionary distance between *S. cerevisiae* and *C. neoformans*, we sought to determine if CnVac8 would bind with homologues of ScVac8 binding partners to carry out the same cellular functions in *C. neoformans*. Unexpectedly, BLAST-P and BLAST-PSI searches of the *C. neoformans* proteome for ScVac8 binding partners revealed few homologues for any of the known proteins.(Error! Reference source not found.) This suggest that despite the conserved protein structure of ScVac8 and CnVac8, their functions may diverge through interactions with unique binding partners. Absent of a Vac17 homolog, the adaptor linking ScVac8 to ScMyo2 for vacuole movement and inheritance, may explain the apparently normal vacuole inheritance in *C. neoformans vac8Δ*. This is not surprising if we consider that in other basidiomycetes, such as *Ustilago maydis*, vacuolar movement and inheritance is kinesin-dependent rather than myosin-dependent like in *S. cerevisiae*.(*52*) Thus, identifying the CnVac8 binding partners might reveal novel biology applicable to other basidiomycetes.

The fungal vacuole is a unique organelle with a diverse set of functions including protein degradation, ionic and metabolite storage, osmoregulation, and regulation of intracellular pH. Given that many of these functions are needed for full virulence in other pathogens, and the need to find novel, fungal-specific, drug targets, we next assessed if the *vac8Δ* strains exhibit defects under common cell stressors. The absence of a closely related organelle to the vacuole in mammalians indicates it may be an advantageous target for development of future therapeutics. *vac8Δ* grew similar to the WT at room temperature, under osmotic stress, nitrosative stress, and with the cell wall stressor CFW. They also displayed modest growth defects at high temperature (37°C), high temperature with CO_2_, and cell membrane stress (SDS). Instead, *vac8Δ* showed a significant growth impairment on media supplemented with caffeine, suggesting *vac8Δ* may be involved in a general cell stress pathway. Surprisingly, at 2 mM hydrogen peroxide, *vac8Δ* exhibits reduced growth, but when exposed to higher levels of 4 mM hydrogen peroxide, it does not appear susceptible and display increased growth compared to WT and *pfa4Δ*. (Error! Reference source not found.) Studies of superoxide dismutase (SOD) mutants in *C. neoformans*, that lead to higher intracellular reactive oxygen species or via menadione treatment, show increased vacuolar fragmentation.(*53*) Therefore we hypothesize, as have others, that vacuolar fragmentation increases the surface-to-volume ratio and allows for more iron transporters to bind and increase iron uptake which serves to protect cells from oxidative stress.(*53, 54*) This same hypothesis might apply to the interesting observation that *vac8Δ* mutants are slightly more resistant to fluconazole without any change in susceptibility to amphotericin B. (**Figure 7**) Recent evidence in *C. albicans* suggests that vacuolar sequestration of azoles through the ATP-binding cassette (ABC) transporter Mlt1p contributes to azole resistance.(*55*) Consistently, deletion of Mlt1p in *C. albicans* (or its homolog in *S. cerevisiae* Ybt1p) leads to azole susceptibility.(*55*) Similar to the yeast vacuole response to oxidative stress, vacuole fragmentation may play a beneficial role in responding to challenge from azoles and potentially facilitate increased sequestration. Supporting this idea is the recent finding that in *C. neoformans*, treatment of cells with fluconazole lead to increased vacuolar fragmentation.(*53*)Since previous vacuolar mutants like *vph1Δ*, which encodes a subunit of the vacuole H^+^-ATPase, are defective in capsule production, laccase production, and growth at 37° C, we also assessed these virulence factors in *vac8Δ*. (*12*) Despite aberrant vacuole morphology, *vac8Δ* shows a modest growth defect at high temperature and modest reduction in capsule production. (Error! Reference source not found.) Both laccase and urease production were similar to WT strains demonstrating that some, but not all virulence factors were affected by deletion of VAC8. These phenotypes could be explained by the fact that although the vacuole is fragmented in *vac8Δ*, it may still be functional, while in the *vph1Δ*, the vacuole is not functional.(*12*) While examining the ability of *vac8Δ* to produce capsule we noted an increase in aberrant cell morphology, including multi-budded cells, and cells with small, elongated buds. We sought to determine if the number of partially formed buds increased over time as that may indicate issues with cell cycle regulation, possibly due to defective vacuole migration into daughter cells.(*56*) In *S. cerevisiae*, the absence of vacuole migration from mother to daughter cell is not lethal under normal conditions because the daughter cell can generate a new vacuole “*de novo*” through unknown mechanisms.(*29*) However, it still appears a functional vacuole is required for cell-cycle progression out of G1 phase, thus any defects in vacuole migration or function may result in cell cycle arrest.(*56*) This migration is an actin-dependent process using the molecular motor Myo2, Vac17, and Vac8 complex.(*29*) Given that we did not find a homologue of Vac17 we were uncertain if Vac8 would be involved in migration of the vacuole and would produce delay in growth. The number of cells displaying abnormal budding did not increase significantly over time in wildtype or *vac8Δ*. However, *vac8Δ* showed much higher levels of incomplete cytokinesis and abnormal bud morphology with very few WT cells having more than one daughter cell (**Figure 5**). Despite not increasing over time, it is clear *VAC8* and normal vacuolar function are required for normal budding behavior.

Along this same line, vacuole size helps control cell size in yeast, initiation of bud formation, and this may also play a role in titan cell formation.(*40, 56*) Using previously establish *in vitro* methods we induced titan cell formation and quantified cell body, capsule, and total cell size. (**Figure 6**) While *vac8Δ* and WT strains were able to produce robust capsule, *vac8Δ* showed severely limited ability to form cells >10 μm.

Examining the vacuole in these titan cells using FM4-64 and DIC highlighted the very large vacuole in the WT cells with a single vacuole encompassing nearly all the internal space. In contrast, *vac8Δ* cells often have multiple vacuoles that do not encompass large portions of the cell body. In some cases, there is one larger vacuole with one or more smaller vacuoles. Together this suggests *VAC8* plays a role in facilitating vacuole- vacuole fusion and/or vacuole enlargement, and the impairment of these processes limits cell body enlargement into titan cells. Given that titanization is also related to cell cycle,(*57*) it is not surprising that under these conditions we also see the abnormal budding morphology in *vac8Δ* mutants, supporting the idea that in *C. neoformans* there is also a vacuolar cell cycle checkpoint.

Lastly, given the cellular and growth defects under certain conditions related to virulence, it was surprising that we found that Vac8 function is not necessary for normal disease manifestation in a murine model of cryptococcosis. (**Figure 8**, **Figure 9**) This may be partially explained by *vac8Δ*’s increased resistance to oxidative stress, which the yeast is likely to encounter in a host, but also potentially to a less restrictive environment in the waxworm relative to mammalian hosts. Also, we hypothesize the absence of *VAC8* leads to increased vacuole fragmentation but does not impair overall vacuolar function. This may explain the increased resistance to oxidative stress and fluconazole and slight growth defects while remaining capable of causing disease. This shows CnVac8 plays a similar role to ScVac8 in regard to membrane tethering and homotypic vacuolar fusion, but also new roles in cell morphology, leading to abnormalities in cell budding and titanization. Because of these unique functions, *VAC8* might still be needed for normal progression and disease manifestation in a mammalian host, similar to how in *C. albicans* vacuole enlargement is needed to promote invasive hyphal growth.(*58*)

Uncovering a role for *VAC8* in *C. neoformans* biology has highlighted an important role of the fungal vacuole in cellular responses to various stresses, including exposure to antifungals and host environments. *ScVAC8*, in particular, has been extensively studied in model yeast because in there, it is the only ARM repeat containing protein. ARM repeats are conserved in eukaryotes, but with distinct and overlapping cellular roles. This diverse list of roles is because ARM repeat containing proteins can bind multiple partners, contributing to multiple functions. Future research on *C. neoformans VAC8* will unveil some of these additional functions; however, through a better understanding of the role of *VAC8* in cryptococcal vacuole we ultimately hope to identify processes and potential targets that may make these fungal pathogens more susceptible to antifungal therapies while also uncovering novel cell biology.

**Table 1:**
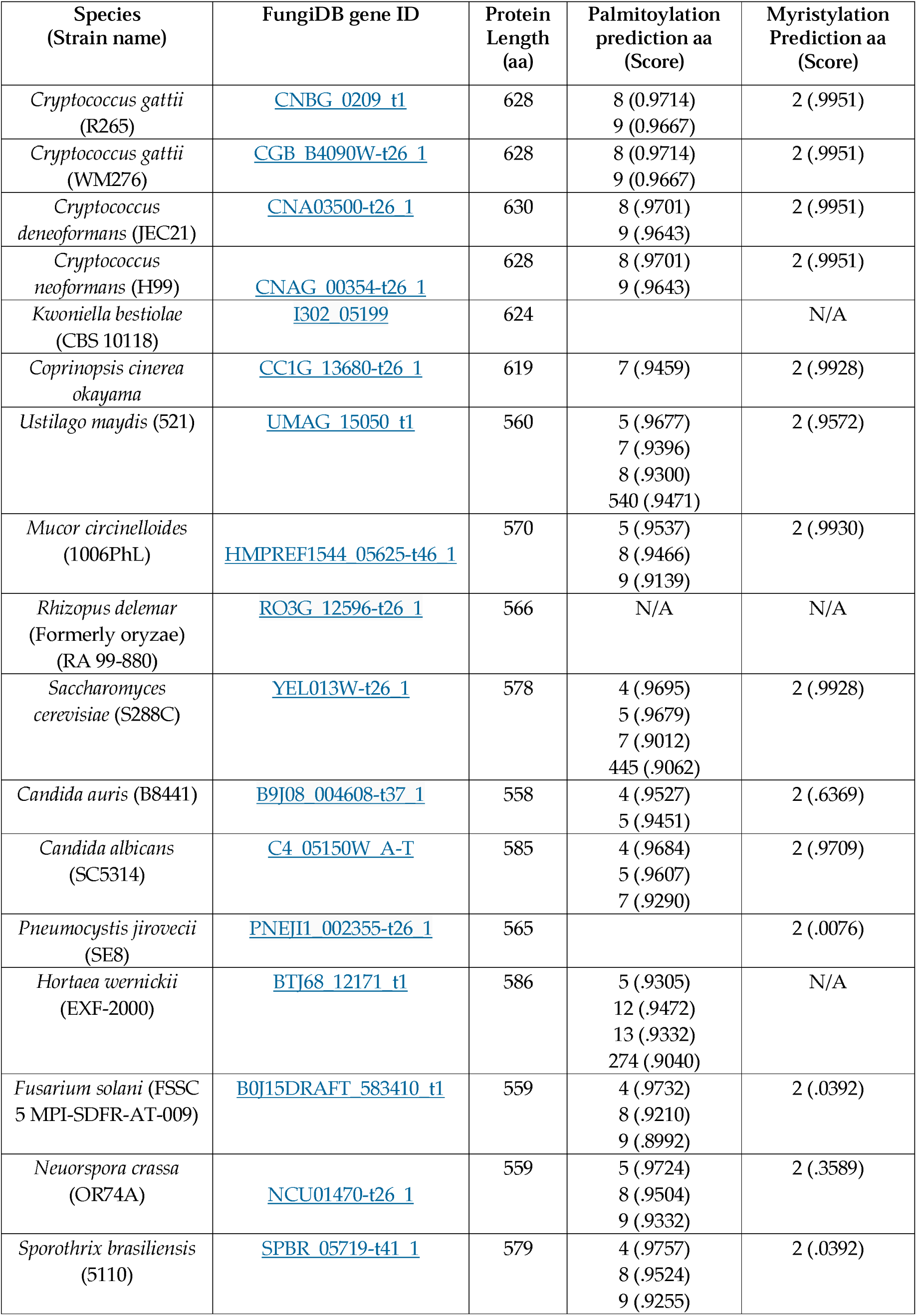
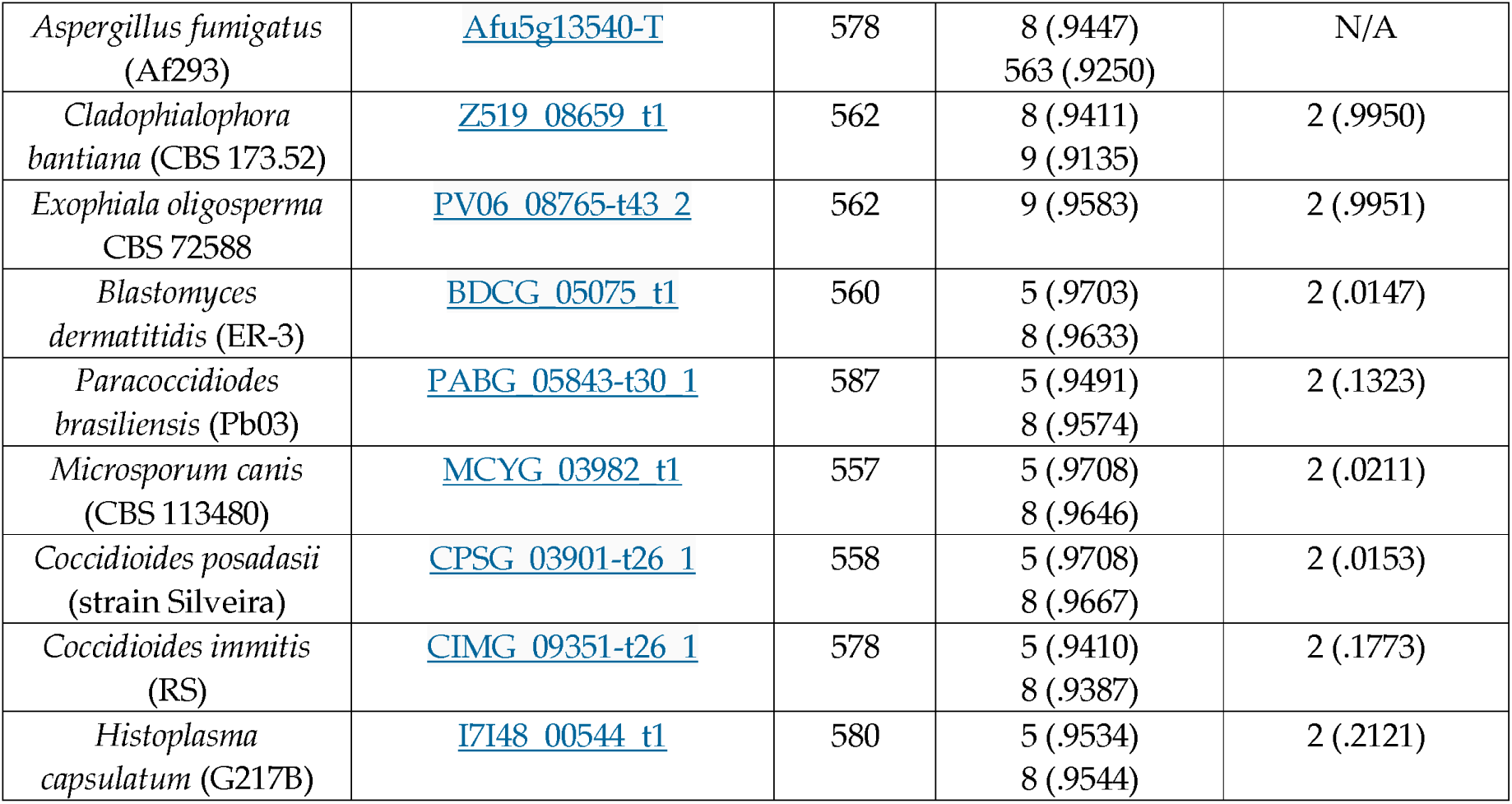
Vac8 protein modification predictions. Protein length and predicted lipid modification with amino acid residues. Confidence threshold of 0.9000 was used as a positive indicator for both palmitoylation and myristylation.(*24, 25*) N/A: there is no N-terminal glycine residue for myristylation.

**Table 1:** Analysis of Vac8 binding partners in *C. neoformans*. *Atg13 has a potential homologue in *C. neoformans* strain H99 (CNAG_00778) with an E value of 5e-4. An E value threshold of 1e-5 was used as a positive indicator of homology for this table.

| Protein Name | Cellular function | Present in *C. neoformans* | E-Value | Ref. |
| --- | --- | --- | --- | --- |
| Nvj1p (YHR195W) | Piecemeal nuclear autophagy | No |  | (60, 61) |
| Vac17 (YCL063W) | For binding to Myo2 complex for vacuole inheritance | No |  | (60) |
| Atg13 (YPR185W) | Required for Cvt pathway and autophagy | No* |  | (60) |
| TCO89 (YPL180W) | TORC complex subunit | No |  | (31) |
| VID21 (YDR359C) | NuA4 histone acetyltransferase complex subunit | Yes<br>CNAG_01591 | 7e-10 | (31) |
| VAB2<br>(YEL005C) | Vac8p binding | No |  | (31, 62) |
| TAO3<br>(YIL129C) | Involved in cell morphogenesis and proliferation | Yes<br>CNAG_03622 | 1e-79 | (31) |

## ACKNOWLEDGMENTS

We would like to acknowledge the minor but important contributions to this project by other undergraduate researchers in the lab, including Rylie O’Meara and Rachel Bartnett. We also thank the other members in the Santiago-Tirado lab that generously served as mentors for these undergraduates, specially PVS, who over the years has been the constant figure supervising this project. This project was possible thanks to institutional funds from the University of Notre Dame. PVS was partially supported by a Paul F. Ware, M.D. Graduate Fellowship, and we acknowledge financial support to MF and AC provided by the College of Science Summer Undergraduate Research Program and BIOS Research Experiences for Undergraduate Program, respectively. Research in the lab of FHST is supported by grants from the National Institutes of Health (R01AI177875 and R21AI171742).

